# The GIAB genomic stratifications resource for human reference genomes

**DOI:** 10.1101/2023.10.27.563846

**Authors:** Nathan Dwarshuis, Divya Kalra, Jennifer McDaniel, Philippe Sanio, Pilar Alvarez Jerez, Bharati Jadhav, Wenyu (Eddy) Huang, Rajarshi Mondal, Ben Busby, Nathan D. Olson, Fritz J Sedlazeck, Justin Wagner, Sina Majidian, Justin M. Zook

## Abstract

Stratification of the genome into different genomic contexts is useful when developing bioinformatics software like variant callers, to assess performance in difficult regions in the human genome. Here we describe a set of genomic stratifications for the human reference genomes GRCh37, GRCh38, and T2T-CHM13v2.0. Generating stratifications for the new complete CHM13 reference genome is critical to understanding improvements in variant caller performance when using this new complete reference. The GIAB stratifications can be used when benchmarking variant calls to analyze difficult regions of the human genome in a standardized way. Here we present stratifications in the CHM13 genome in comparison to GRCh37 and GRCh38, highlighting expansions in hard-to-map and GC-rich stratifications which provide useful insight for accuracy of variants in these newly-added regions. To evaluate the reliability and utility of the new stratifications, we used the stratifications of the three references to assess accuracy of variant calls in diverse, challenging genomic regions. The means to generate these stratifications are available as a snakemake pipeline at https://github.com/ndwarshuis/giab-stratifications.

## Introduction

The Genome in a Bottle (GIAB) consortium generates variant benchmarks for a set of human genomes to enable evaluation and comparison of sequencing technologies and variant detection methods^1–3^. GIAB has expanded its variant calling benchmark sets to include increasingly challenging genomic regions and variants as sequencing technologies, variant detection methods (for both single nucleotide variants (SNVs) and structural variants), and assembly algorithms improve ^4^. Current technologies can resolve variants in most of the genome, but correctly calling variants in complex or repetitive regions remains a challenge ^5,6^. GIAB continues to build a metrology infrastructure for assessing performance of variant detection methods, including files denoting different regions of the genome, which we term stratifications, that can be used to identify strengths and biases in genome sequencing and variant calling ^7,8^. Coding regions, low mappability regions, high GC content regions, and various types of repetitive regions are examples of such genomic stratifications (we describe all stratification categories in more detail in Supplemental Note 1). The challenges here are multifold to ensure that these stratifications are harmonized across different genomes.

Genomic stratifications provide value for many users in the genomics community, including bioinformatics method developers, sequencing technology developers, and clinical laboratories. For those developing software tools, stratifications can be used to better understand the advancements or limitations of new methods and identify biases across methods or technologies. For example, stratifications were used in the precisionFDA challenge V2 to evaluate the performance of different technologies across different repetitive regions, such as homopolymers or segmental duplications^8^. Additionally, with respect to evaluating the performance of calling insertions and deletions (INDELs), this study revealed that the INDEL recall and precision metrics are lower when using PCR amplification compared to PCR-free sequencing calculated for the whole genome. However, these values are almost equal when considering all regions except homopolymers or tandem repeats. This observation demonstrates the importance of genomic stratifications and how they can highlight differences between different technologies and pipelines, allowing for critical investigation^7^. This information can help end users make informed trade-offs when selecting tools, where performance vs runtime, server cost, hardware requirements, reagent costs, and user expertise must be balanced.

Stratifications are also important in medical practice, both at the research level and in the clinic. For the researcher, stratifications indicate genomic regions where “difficult” variants might be found and as such might require additional resources to study accurately. Stratifications also carry some functional and/or structural information, such as specifying which regions contain coding genes^9^ or high GC content, which is useful for designing experiments and association studies. For the clinician, stratifications provide a means to assess confidence in a result. Guidelines for validating clinical pipelines include validating “representative” variants of different types and genome contexts, and stratifications define genome contexts that are challenging^10^. If a patient presents with a pathogenic variant, stratifications can show if this variant resides within a “difficult” region, which in turn could provide a proxy for how much the clinician can trust the result. Thus, stratifications are instrumental in the development and understanding of variants across different disciplines.

Given the multifaceted importance of stratifications, their continuous improvements are essential. Stratifications are currently defined with regard to two linear references, GRCh37 and GRCh38. The genomic stratifications were originally developed in collaboration with the Global Alliance for Genomics and Health (GA4GH)^7^ and are being maintained by GIAB. As part of this study we developed a set of comparable stratifications for the CHM13 human genome reference. CHM13 was recently published as the first Telomere-to-Telomere (T2T) reference^11^ which completed the remaining 8% of gaps present in the existing references, adding ∼2000 genes and ∼100 protein coding sequences. In other words, CHM13 provides gapless assemblies for all chromosomes by introducing about 200 Mbp covering both euchromatic and heterochromatic regions. Furthermore, it includes centromeric satellite arrays, segmental duplications, and the short arms of all five acrocentric chromosomes^11,12^. Overall CHM13 has shown to improve sequencing data analysis including variant calling^11^. To fully leverage such a reference genome and assess the reliability of existing methods, a set of genomic stratifications for CHM13 is needed. This facilitates the study of hundreds of new genes and their role in phenotypes or diseases. Moreover, developers of sequencing technology and genome assessment pipelines can benefit from CHM13 and associated stratifications in understanding performance in difficult, newly assembled regions.

Stratifications could also be expanded to include new types of genomic context that offer more nuanced information regarding the expected error mechanism. For example, it is much easier to call a variant in a tandem repeat if it is the only variant in the repeat; additional variants could “shift” the representation of the variant being called which makes variant calling and variant comparison challenging. Similarly, current stratifications do not account for read coverage or distance between variants. The former is important as higher coverage may imply less difficulty. The latter hinders variant calling where variants are closer together probably due to representational challenges. Accordingly, the current study examines the possible benefits from the inclusion of new stratifications characterizing read coverage and variant distributions.

## Results

The GIAB stratification resource is a publicly available dataset for the human reference genome. Here we describe the extension of this resource to the CHM13 reference genome (Table 1). We also provide an insight to the differences between three reference genomes: GRCh37, GRCh38 and CHM13. Furthermore, we explore three new features for future stratifications of GRCh38 which include variant distributions in tandem repeats, read coverage of variants, and genomic distance between variants. We now automated generating all stratifications to further facilitate the creation across upcoming complete human genomes or other reference genomes: https://github.com/ndwarshuis/giab-strati?cations.

**Table 1:** Overview of existing and newly-added stratification types for the three reference genomes. Blue rows are new stratification categories that might be added to future versions. Checkmarks denote existing stratifications. “X” denotes stratifications that were not produced previously for a given reference. “This study” denotes stratifications that were added in this work. Stratifications denoted with NA are not covered in this study.

| **Stratifications** | **GRCh37** | **GRCh38** | **CHM13** |
| --- | --- | --- | --- |
| Gene coding regions | ✓ | ✓ | This study |
| Functional, technically difficult to sequence | ✓ | ✓ | NA |
| GC content | ✓ | ✓ | This study |
| Genome specific | ✓ | ✓ | NA |
| Regions with low complexity sequences | ✓ | ✓ | This study |
| Other difficult genomic regions | ✓ | ✓ | This study |
| Segmental duplications | ✓ | ✓ | This study |
| Chromosome XY specific regions | ✓ | ✓ | This study |
| Patterns of local ancestry | X | ✓ | NA |
| Low mappability regions | ✓ | ✓ | This study |
| Variant complexity in tandem repeats |  | This study |  |
| Distribution of genomic distance between consecutive variants |  | This study |  |
| Read coverage of each variant |  | This study |  |

### Extending CHM13 stratifications

#### Gene coding regions

Coding sequences (CDS) are the regions of the genome that code for proteins which are usually targeted for many clinical tests^4^. Using the CHM13v2.0 assembly and available RefSeq annotations^9^, we were able to extract gene coding regions and compare them with those of GRCh38 and GRCh37. The methods used for generating CDS stratifications for GRCh38 were applied to CHM13v2.0^8^. Figure 1 shows the comparison between the total length of the CDS region and its ratio over chromosome length (excluding unknown bases in the assembly which are noted with N) for the all three references, GRCh37, GRCh38 and CHM13. All three reference genomes have similar CDS coverage across chromosomes. If the CHM13 CDS annotation was modeled after GRCh38 annotations, we would expect similar results. Also, the differences that we see in chromosomes 9, 19, and 22 are likely due to the fact that several new bases were added in CHM13 relative to GRCh38 on these chromosomes as they have large centromeres and heterochromatin that were excluded in GRCh38^11^.

**Figure 1:**
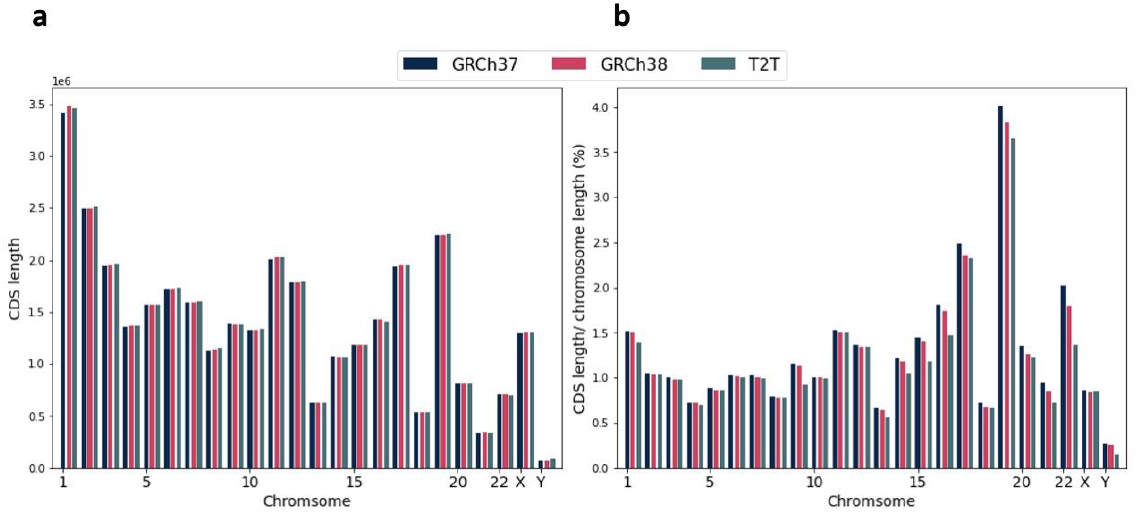
**(a)** Total length of CDS regions for GRCh37, GRCh38 and CHM13. **(b)** The ratio of length of CDS regions over chromosome length (excluding unknown bases) for GRCh37, GRCh38 and CHM13.

#### Low mappability regions

Reference mappability is a metric that can be used to identify whether reads of a given length will align uniquely to that region of the genome (lower means to harder-to-map). The regions that are found to be of low mappability had been previously generated for GRCh37 and GRCh38. We calculated such regions for the CHM13 reference genome at two different stringencies. For moderately difficult-to-map regions we permitted up to two mismatches and one INDEL between each 100 bp region and any other region. However, for very difficult-to-map regions we permitted no mismatches or INDELs between each 250 bp region and any other region. The total length of low mappability regions for each chromosome is depicted in Figure 2 for all three reference genomes. The total length of the regions in the CHM13 is higher than that of the older references, particularly for chromosomes 1 and 9 which have large satellite repeats. In addition, the large increases in low mappability regions in CHM13 relative to the previous references can be partly explained by the addition of rRNA repeat arrays on the acrocentric chromosome arms (13, 14, 15, 21, and 22)^12,13^. Overall, the low-mappable regions vary among different chromosomes and reference genomes.

**Figure 2:**
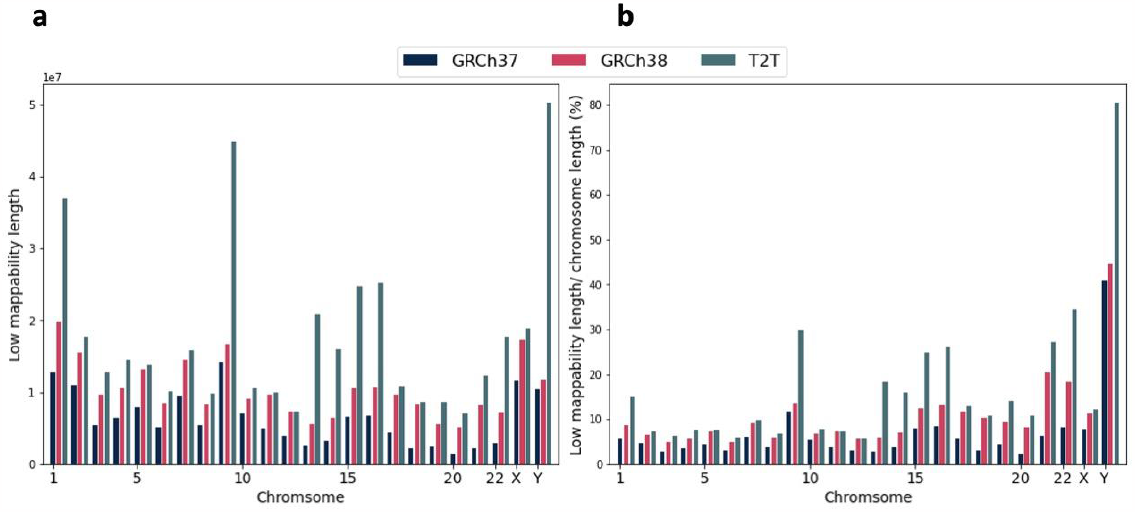
Statistics of low-mappable regions in GRCh37, GRCh38 and CHM13. **(a)** Total length of low-mappable regions, and **(b)** Ratio of total length over chromosome length (excluding unknown bases) for GRCh37, GRCh38 and CHM13.

#### GC content

Defining high and low GC-content regions (i.e., the fraction of G and C bases is high or low) is important as different sequencing technologies can produce distinct error profiles in GC-rich and AT-rich regions^14^. This stratification delineates regions with specific amounts of GC content. These are based on the method used for generating standardized GC content browser extensible data (BED) files by the Global Alliance for Genomics and Health (GA4GH) Benchmarking Team and the Genome in a Bottle (GIAB). Of note, we consider 10 different ranges from 15 to 85% of GC content with interval length of 5% in addition to two cases of regions with GC content of smaller than 15% and greater than 85%.

As an example, we depicted the total length of regions for each chromosome of the human reference genomes GRCh37, GRCh38 and CHM13 with the GC content in the range of 20-25% in Figure 3a. The ratio of total length of regions over chromosome length (excluding unknown bases) is illustrated in Figure 3b and 3d. As we can see, three reference genomes follow a similar pattern except for chromosome 13 in CHM13. Moreover, Figure 3c shows the total length of regions with GC content higher than 85% for GRCh37, GRCh38 and CHM13. It is evident that the values for chromosomes 13, 15 and 21 are much higher in CHM13 compared to other references. These could be explained by considering the fact that these are the human acrocentric chromosomes which have cytogenetically similar short arms^13^ and there is a potential to investigate this with further analysis.

**Figure 3:**
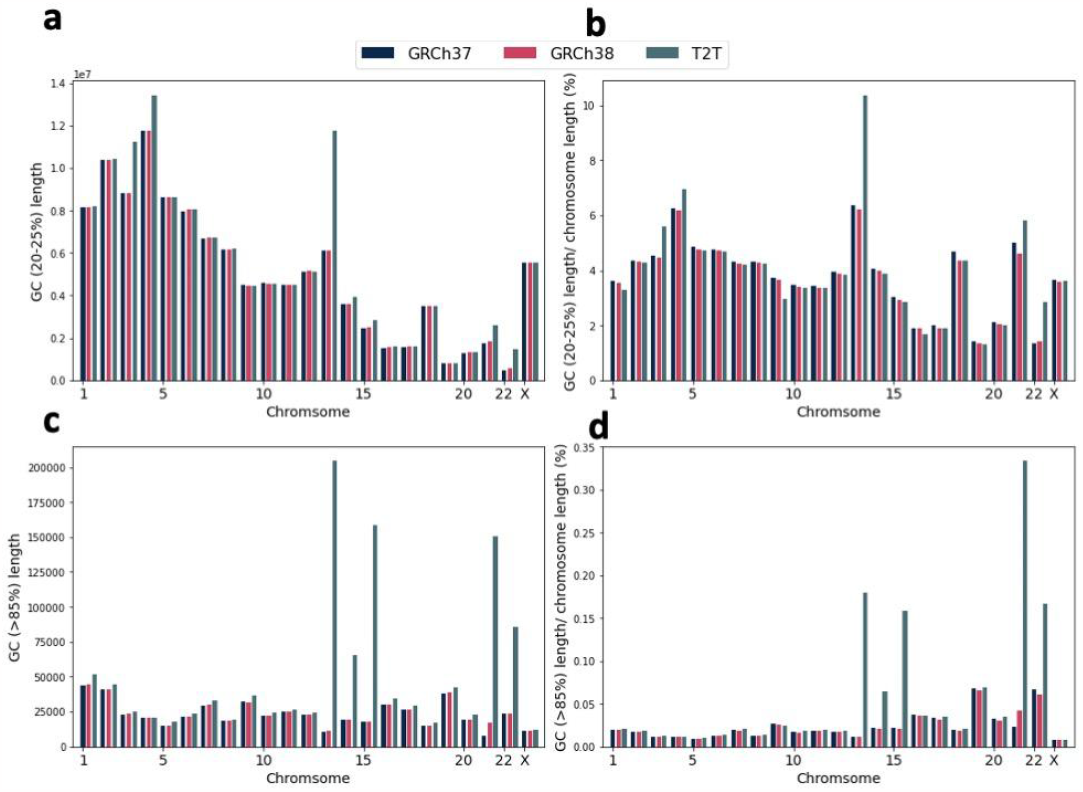
Statistics of regions with specific GC content for GRCh37, GRCh38 and CHM13 **(a)** Total length of regions with GC content in the range of 20-25%. **(b)** Ratio of total length of regions with 20-25% GC content over chromosome length (excluding unknown bases). **(c)** Length of regions with GC content higher than 85%. **(d)** Ratio of total length of regions with GC content >85% over chromosome length (excluding unknown bases).

#### Other challenging, medically-relevant genomic regions

Three challenging, medically-relevant regions within human genomes including the Major Histocompatibility Complex (MHC), variable/diversity/joining (VDJ) and Killer-cell immunoglobulin-like receptor (KIR) are considered here^15^. These three regions are all highly polymorphic and underpin key immunological functions: the MHC region contains the Human Leukocyte Antigen (HLA) genes which determine “donor matches”, the VDJ regions are randomly recombined to produce the T and B cell receptors, and the KIR region codes for one of the key effector receptors on natural killer cells. The total length of each region on the three reference genomes, GRCh37, GRCh38 and CHM13, are reported in Table 2. As shown, the regions are located on the same chromosome across different reference genomes with comparable total length.

**Table 2.** Total length of three difficult genomic regions, namely MHC, VDJ and KIR, on reference genomes GRCh37, GRCh38 and CHM13.

|  | Chromosome(s) | GRCh37 | GRCh38 | CHM13 |
| --- | --- | --- | --- | --- |
| **MHC** | 6 | 4,970,558 | 4,970,558 | 4,920,493 |
| **VDJ** | 21,422 | 3,232,649 | 3,348,726 | 3,541,519 |
| **KIR** | 19 | 155,001 | 155,044 | 156,419 |

### Benchmarking new stratifications

We evaluated the new stratifications for their most common use, which is benchmarking performance of methods in different types of difficult regions. Regions unique to CHM13 relative to GRCh37/38 affected benchmarking performance for certain variant types. For example, a much larger fraction of CHM13 is satellite tandem repeats due to the newly assembled centromeres. We benchmarked variant calls on each reference using hap.py (see Methods) with an example long read-based callset (HiFi-DeepVariant) of HG002 sample as the query and a draft benchmark based on the HG002 assembly from the Human Pangenome Reference Consortium (HPRC)^16^ as the truth. The draft benchmark excludes regions not covered by the assembly and partially covered repeats so that most but not all assembly errors are excluded.

To assess the reliability of stratifications, we compared performance metrics for the new stratifications between references to ensure that metrics are similar or we can explain the differences (Supplemental File 1). In general, precision and recall were comparable between references and highest for non-difficult regions as expected (e.g. *notinalldifficult*) which was >99% for SNVs, and close to 99% for INDELs. CHM13 had low precision/recall (50% or less) in the chromosome X Pseudoautosomal regions (PAR) because the Y PAR region was not masked when aligning reads to CHM13, showing the utility of stratifying the PAR region for diagnosing problems. CHM13 also had a comparable or higher number of not assessed variants (i.e., variants that fell outside the confident benchmark regions), particularly for SNVs, INDELs, and large deletions. This was likely due to the considerably higher amount of sequence added to CHM13 that contains satellites and segmental duplications with complex structural variation. Precision and recall in segmental duplications were lower in CHM13 for all variant types, which is expected given the larger number of segmental duplications included in CHM13. Our new stratifications separating A/T and G/C homopolymers show higher error rates for G/C homopolymers than A/T homopolymers.

### Exploring new features for future stratifications

In this study we expanded several of the stratifications to CHM13. We also explored possible new stratifications for GRCh38 in addition to the so far described well-established stratifications. We examined variant complexity in tandem repeats, read coverage for each variant with a given bam file, and the genomic distance between variants in the whole genome.

#### Exploring variant complexity in tandem repeats

Tandem repeats frequently contain complex variants (e.g., multiple SNVs and INDELs), which can cause errors in variant calling and benchmarking. Variant callers tend to produce more errors in repeats because variants could be mis-identified among the repeat, particularly if reads are insufficiently long to contain the entire repeat. Notably, such errors are likely to increase when there is more than one variant in the region. It may thus be sensible to create new stratifications for tandem repeats categories by number of variants, where number of variants corresponds to difficulty.

To understand how variants were distributed in tandem repeats, we intersected the variant call format file (VCF) derived from an HPRC assembly of the HG002 sample^16^ with the GIAB v3.1 tandem repeat and homopolymer stratification BED files. This VCF includes both small variants and structural variants (SVs) (except for inversions and translocations), so that we can assess the full spectrum of variants in tandem repeats^2^. After splitting multiallelic variants, we additionally filtered any variants that overlapped repeat boundaries (∼1500 variants), though these would be important sources of complexity to explore in the future. We found that the vast majority (∼90%) of the repeats in GRCh38 did not have any variation, but >10,000 tandem repeats contain more than one variant and >1,000 contain more than three variants, resulting in complex variants that can cause challenges in variant calling and variant representation (Figure 4a).

**Figure 4:**
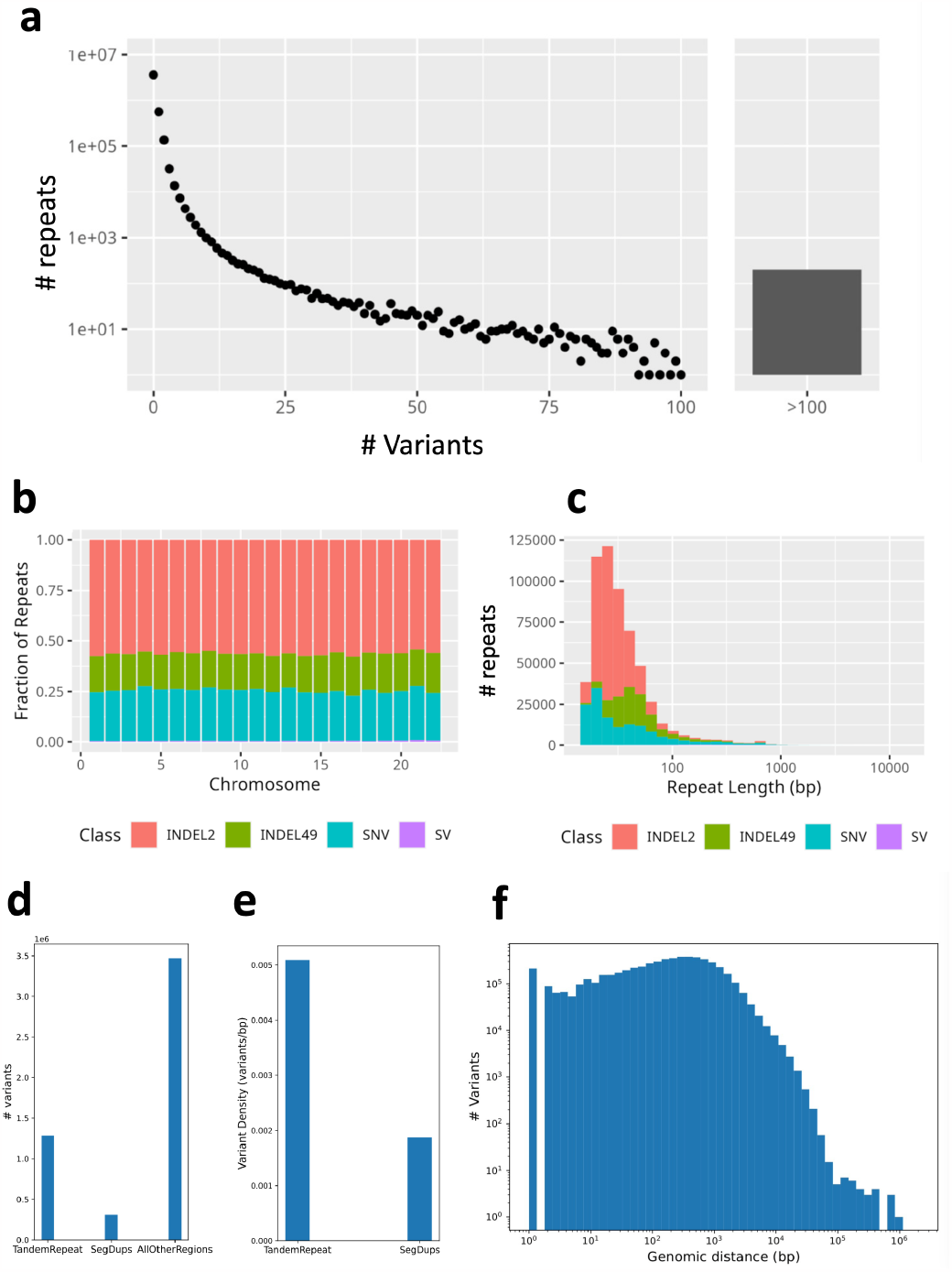
Distribution of SNV and small INDEL variants within tandem repeats throughout GRCh38 using a diploid assembly-based VCF from HPRC. **(a)** The distribution of number of variants per repeat. Y-axis shows the number of tandem repeats and x-axis is the number of variants in each tandem repeat. **(b-c)** Among repeats with only one variant, the fraction of the variant class by chromosome **(b)** and the distribution of intersecting variants classified by type according to repeat length **(c)** INDEL2; INDELs with length <= 2, INDEL49; INDELs with length > 2 and length <= 49, SNV; single nucleotide variants, SV; structural variants. **(d)** Number of variants in tandem repeats, segmental duplications and all other regions. **(e)** Variant density in regions of tandem repeats and segmental duplications. **(f)** The distributions of genomic distance between any two consecutive variants for all autosomal chromosomes for HG002.

We further investigated the size distribution of repeats with single variants. About 30% of such variants were SNVs, and about 50% were INDELs between 1-2 bp (Figure 4b). There were several hundred SVs associated with these repeats as well, indicating that some SVs in repeat regions exist without any simpler variants in the same repeat. This distribution was mostly consistent across chromosomes. Furthermore, we investigated the size distribution of repeats with a single variant according to variant type (Figure 4c). We found that small INDELs are disproportionately more likely to be in repeats < 80 bp long compared to other variant types. Of note, the variant density in tandem repeats is more than two times of that in segmental duplications (Figure 4d-e).

#### Exploring distribution of genomic distance between consecutive variants

Understanding variant distribution is important, since the likelihood of representational difficulties increases as the distance between any two variants decreases. Furthermore, in extreme cases, too many variants within a region can lead to a reference allele bias due to e.g. mapping biases^17,18^. Conversely, read-based phasing of variants becomes more challenging as distance between heterozygous variants increases^6,19,20^.

Here we analyze the variants for sample HG002 when aligning a diploid assembly from HPRC to GRCh38, which captures a broad range of variant types with less reference bias. We depicted the distribution of the genomic distance between any two consecutive variants for all autosomal chromosomes (Figure 4f). The furthest neighboring variants appeared in the centromere of chromosome 12 starting at position 35,094,141 with a distance of 2,018,782bp. The next three largest distances are in the centromere of chromosome X and 17. Of note, 2.1 million (38.6%) of the 5.5 million variants have a neighboring variant within 100bp. Interestingly, more than one-third of these neighboring variants are in repetitive regions including tandem repeats and segmental duplications, which may cause variant calling and representation challenges. Notably, chromosome 1 encompasses 424,478 variants with a median genomic distance of 211 bp. Additionally, 151,165 of the variants (35.6%) have a distance of <100bp and the number of neighboring variants with only one base afar is 13,264 (Supplementary Figure 1).

#### Exploring read coverage of each variant

In addition to stratifications mentioned above, here we consider the HG002 individual to explore the read coverage of each variant, where abnormal coverage implies that the region flanking the variant is a difficult region to align or call variants due to potential misrepresentations (e.g. copy number variants). Additionally, the idea behind calculating the coverage values is to look for new informative metrics, which can be fed as an additional feature into machine learning models to predict the quality of called variants^21^. Obviously, one would expect with higher coverage the variants to be more robustly identified. However, too high coverage may indicate mis-mapping of reads due to duplications in the individual, which can result in false variant calls. Another cause of abnormal coverage is small deletions and insertions, so we have chosen to look at three different variant types: deletion (DEL), insertion (INS) and single nucleotide variant (SNV). This approach could be explored by calculating the read coverage values of the BAM file in the positions of these three variant types (DEL, INS and SNV) called using dipcall with HPRC diploid assembly of GIAB HG002.

Accordingly, we calculated the average read coverage of genomic positions in each variant type (see Methods). The average coverage values of DEL, INS and SNV positions are 38.2, 49.9, and 58.8, correspondingly, which shows their unique characteristics. Of note, the average coverage of positions in deletions (38.2) are much lower than that of SNVs (58.8) which is expected due to homozygous or heterozygous deletions. However, this needs further investigations to evaluate variant quality.

## Discussion and Conclusion

In this work we summarized and reported a complete set of genomic stratifications across GRCh37, GRCh38 and CHM13. We showcased multiple unique properties of these stratifications and give in depth explanations about their importance and relevance for the genomic community. Thus, we are confident that these stratifications will be leveraged across multiple experiments and method development projects to obtain deeper insights. In addition to previously used stratifications we now introduce several new ones that should yield even further insights into different methods. As such we introduce the variant distance that reflects the complexity of a region, the variant coverage that will provide deeper insights into biases due to too high or too low coverage regions (e.g. due to repetitive regions or reduced sequencing performance). This work represents a rosetta stone to better understand variant analysis and will be utilized across consortia and single sample projects that rely on standardized stratifications to filter and optimize their methodologies. This presented resource will be beneficial for several other applications even outside of GIAB. For example, the stratifications of regions with low complexity sequences could be used to filter variants in repeat regions^22^ and abnormal coverage values could be utilized for quality control of haplotype phasings^23^. Also, such features have the potential to be used for improving and filtering gene annotations^24^, gene expression^25^, genome-wide association studies^26^. Finally, our pipeline will enable stratifications to be generated for additional assemblies, such as the T2T diploid assembly for HG002^27^, which can be used to flag difficult regions which in turn will enable easier constructure of future human diploid assemblies.

GIAB continues maintaining the current stratifications and generating new ones. Specifically, additional stratifications could be generated for additional pangenome references in the future, such as those being developed by the Human Pangenome Reference Consortium. In this paper, we reviewed 10 stratifications of the human reference genomes GRCh37, GRCh38, and CHM13 and compared four of them including regions of low mappability, high GC content, CDS and other difficult genomic regions. We also explored three new genomic stratifications including variant complexity in tandem repeats, variant read coverage, and distribution of genomic distance between consecutive variants. Future genome-specific stratifications could include these regions to assess how variant complexity influences accuracy.

## Methods

### Exploring mappability of CHM13 using GEM

We used GEnome Multitool (GEM)-Mapper ^28^ (version pre-release 3) on the CHM13v2.0 reference genome to create BED files of low mappable regions. We followed available scripts https://github.com/genome-in-a-bottle/genome-stratifications/tree/master/GRCh38/mappability. Briefly, we generated raw mappability files under two stringency levels: low stringency (100 bp single-end reads, two mismatches, and one INDEL) and high stringency (250bp single-end reads, 0 mismatches, and 0 INDELs). These mappability BED files were then processed with SAMtools and BEDtools to find the union of the two stringency levels for the final BED file with low mappability regions for CHM13v2.0.

After running both mappability scripts, we had four final BED files containing non-uniquely mapped regions for each stringency level, as well as a final BED file containing all low mappability regions when performing the union for both stringencies.

### Generate CDS regions for CHM13v2.0

We used the R script provided for GRCh38 https://github.com/genome-in-a-bottle/genome-stratifications/blob/master/GRCh38/FunctionalRegions/create_GRCh38_cds_bed.Rmd and ported it over to identify the gene coding regions (CDS) in the CHM13v2.0 assembly. It requires R packages - rmarkdown, tinytex, knitr, tidyverse, devtools.

We used the following files from the NCBI FTP site:

- FTBL: ftp://ftp.ncbi.nlm.nih.gov//genomes/refseq/vertebrate_mammalian/Homo_sapiens/all_assembly_versions/GCF_009914755.1_T2T-CHM13v2.0/GCF_009914755.1_T2T-CHM13v2.0_feature_table.txt.gz
- GFF: ftp://ftp.ncbi.nlm.nih.gov//genomes/refseq/vertebrate_mammalian/Homo_sapiens/all_assembly_versions/GCF_009914755.1_T2T-CHM13v2.0/GCF_009914755.1_T2T-CHM13v2.0_feature_table.txt.gz

Additionally, the script required a .fai index file which was created from the CHM13v2.0 reference assembly.

### Generating GC content BED files using seqtk for CHM13v2.0

We use an existing script created to generate the GRCh38 GC Content Stratification BED files. The script required seqtk version-1.3-r106 tool, bedtools v2.27.1, and tabix v1.9. Three essential data files were required to run the script file: the CHM13v2.0 FASTA, the CHM13 genome file. The genome was converted to BED format by adding a middle column of 0 (such that each line had the length of the entire chromosome). We ran seqtk for various fractions of GC content, all within windows of 100bp. After running seqtk, we added 50bp slop to each BED file and merged.

### Lift-over for OtherDifficult regions

In order to find the coordinate of well-studied genes including MHC, KIR, and VDJ that are considered as difficult regions, we performed liftover for such regions from GRCh38 to CHM13v2.0. To obtain the OtherDifficult regions data of the GRCh38 we referred to the reference sample released by the GIAB https://ftp-trace.ncbi.nlm.nih.gov/ReferenceSamples/giab/release/genome-stratifications/v3.1/GRCh38/OtherDifficult/. To perform the lift-over, we used the minimap2 (v2.24) aligner with arguments -ax asm5 followed by bedtools bamtobed and merge (v2.30.0) . The resulting BED files are provided as part of the GIAB stratification resource.

### Exploring variant distributions in tandem repeats

The GIAB HG002 v4.2.1 VCF was converted into a BED-like format where the chromosome coordinates were defined using the POS column and the length of the REF field. The length of the variant (length(ALT) - length(REF)) was also stored. This BED-like file was then intersected (left outer join) with the Tandem Repeats/Homopolymers stratification BED file from GIAB v3.1 stratifications. We ignored multiallelic variants as well as variants that were partially outside a repeat region to simplify analysis.

### Exploring read coverage of each variant

We used the called variants for the HPRC diploid assembly of GIAB HG002 aligned to GRCh38 with variants called using dipcall https://ftp-trace.ncbi.nlm.nih.gov/ReferenceSamples/giab/data/AshkenazimTrio/analysis/HPRC-HG002.cur.20211005/HPRC-cur.20211005-align2-GRCh38.dip.vcf.gz. We consider three different variant types, deletion (DEL), insertion (INS) and single nucleotide variants (SNV). For the extraction of the coverage, the mosdepth package v0.3.2 was used with the flag of setting the bin size to 1 based on the 40x PCR-free HiSeq X bam file at https://storage.googleapis.com/brain-genomics-public/research/sequencing/grch38/bam/hiseqx/wgs_pcr_free/40x/HG002.hiseqx.pcr-free.40x.dedup.grch38.bam ^29^.

### Exploring distribution of genomic distance between consecutive variants

The genomic positions of variants for the sample GIAB HG002 were extracted from the same diploid assembly VCF used above for coverage of each variant. The distance between two consecutive variants were calculated. Using the matplotlib package v3.4.3 of Python v3.9, the histogram figure was depicted. We should also note that a portion of the genome is unknown (existing as Ns in the reference file), so no variant can be found in these regions. To make sure this fact does not influence our analysis, we discarded those variants neighboring unknown sequences in the reference genome which accounts for 328 variants out of 5,596,945 variants.

### Snakemake pipeline

#### Overview

This work (first done as part of a hackathon) was incorporated into a snakemake pipeline which can be found at https://github.com/ndwarshuis/giab-strats-smk and https://github.com/ndwarshuis/giab-stratifications. The latter repository holds the global configuration for the three references in this work, and references the former repository as a submodule. The former repository is reference-agnostic and encodes the build rules for the stratification files themselves.

For the identity of every input file used to make these stratifications, refer to https://github.com/ndwarshuis/giab-stratifications/blob/master/config/all.yml.

#### Stratification validation

Each stratification file prior to publishing was ensured to meet the following criteria:

- Only contained valid chromosomes (ie 1-22, X, Y)
- File was bgzip compressed
- File was a valid bedfile (three columns, tab-delimited, with 2nd and 3rd columns as non-negative integers with 3rd greater than 2nd)
- All regions in the bed file were sorted in numeric order (ie chromosomes ordered 1-22, X, then Y with each region then sorted by start and end)
- No regions overlapped with each other
- No region overlapped a gap region (which included the PAR on chromosome Y)
- No region fell outside chromosomal boundaries

#### Benchmarking

To show how the stratifications could be used to assess performance, we benchmarked long-read-based callsets with an assembly-based benchmark for each reference. The benchmark was created using polished HPRC assemblies for HG002 and mapped against each reference using dipcall. The output VCF and bed files from this were used as the truth VCF and confident regions respectively.

We ran the benchmark using happy as follows:

`hap.py –engine vcfeval –stratifications <path/to/strats> –f`

`<path/to/confident_regions.bed> -o <path/to/output>`

`<path/to/bench.vcf> <path/to/query.vcf>`

## Data and software availability

All versions of the genome stratifications up to v3.2 (the latest as of this writing) are available on an FTP site hosted by NCBI here at https://ftp-trace.ncbi.nlm.nih.gov/ReferenceSamples/giab/release/genome-stratifications/ and are described at https://github.com/genome-in-a-bottle/genome-stratifications/. The initial work for this study (which originally took place at a hackathon) is freely available https://github.com/collaborativebioinformatics/NIST-GREX. A copy of the GitHub repository and HTML output of the snakemake pipeline are archived at Zenodo at https://doi.org/10.5281/zenodo.8414359.

## Author contributions

N.D., F.J.S, J.W., S.M., and J.M.Z designed the study. N.D. implemented the pipeline. N.D., D.K., J.M., N.D.O, P.S, P.A.J, B.J., E.H., R.M. and S.M. performed the analyses. N.D., B.B.,

F.J.S, S.M., and J.M.Z organized the study. All authors reviewed and approved the manuscript.

## Acknowledgements

We thank Sierra Miller and Katherine Gettings for their feedback. Certain commercial equipment, instruments, or materials are identified to specify adequately experimental conditions or reported results. Such identification does not imply recommendation or endorsement by the National Institute of Standards and Technology, nor does it imply that the equipment, instruments, or materials identified are necessarily the best available for the purpose.

## Supplementary Information

### Supplemental Note 1

#### Stratification overview

In this section, we provide an overview of existing genomic stratifications for GRCh37/38 including regions with high/low GC content, low complexity, low mappability, and segmental duplications, in addition to regions with patterns of local ancestry, chromosome XY specific regions and other difficult genomic regions.

##### Gene coding regions

Coding sequences (CDS) are the regions of the genome that code for proteins. These are usually targeted for many clinical tests. Since many diseases are caused by variants in protein sequences, being able to find variants in their corresponding genome sequences is important in clinical applications^4^. For GRCh37 and GRCh38, the coding regions were defined using the RefSeq annotations dataset^9^.

##### Low mappability regions

To identify whether reads of a given length will align uniquely to a region of the genome, the reference mappability metric is utilized. One method to identify such regions is part of the GEnome Multitool (GEM)^28^. This tool queries the genome for sequences of a given length that align other places; these alignments may be allowed some upper limit of mismatches and gaps, corresponding respectively to SNVs and INDELs. For GRCh37/38, current stratifications include two levels of mappability stringency. The “low stringency” set includes all regions 100bp long that align somewhere else within the genome with no more than two SNVs and one INDEL. The “high stringency” set includes all regions 250bp long that perfectly align other places within the genome without any SNVs or INDELs. The lengths for these mappability stratifications roughly correspond to those commonly provided by short read sequencing technologies.

##### GC Content

The regions where the fraction of G and C bases is high or low are simply defined as high and low GC-content regions. Such definition is important as different sequencing technologies can produce distinct error profiles in GC-rich and AT-rich regions^14^. For example, GC-rich and AT-rich regions tend to have reduced coverage for many technologies^30^. GC-rich and AT-rich regions also can have reduced precision and recall in variant calling^31^. For GRCh37/38, these regions have been determined using the seqtk algorithm^32^. We identify regions with specific GC percentage in 5% increments from 15-85%.

##### Segmental Duplications

DNA sequences longer than 1kb with high sequence identity (typically >90%) that are repeated in a genome are called segmental duplications (segdups)^33^. Segmental duplications play important roles in genome evolution and many diseases^34^. The GIAB stratifications define segdups based on two data sources. The first is defined from genomicSuperDups^35^ hosted at the UCSC genome browser. This dataset defines the “canonical” segmentally duplicated regions, e.g., regions >1kb with 90% similarity without other repeat regions such as LINEs or SINEs. These stratification regions were identified by merging all regions from either source database and removing the Pseudoautosomal regions (PAR) from the X and Y chromosomes because these were incorrectly determined to be segmental duplications. The stratifications include two sets of BED files for each source: one with all regions and one with regions >10kb.

##### Regions with low complexity sequences

Another genomic stratification indicates regions with low complexity sequences. We define “low complexity” to include perfect and imperfect homopolymers, tandem repeats, and satellites. A homopolymer is defined as a sequence of consecutive identical bases. Tandem repeats refer to motifs that repeat up to a given length. Depending on the motif length they can either be short tandem repeats (STRs, also known as microsatellites) which have a motif size of 2-6 bases or variable number tandem repeats (VNTRs) which can have longer motifs. Satellites have repeating motifs like tandem repeats, but the total length of the repeat is generally much longer and motifs are more complex. The longest satellites occur in centromeric regions.

To construct the low complexity stratifications we utilized several sources. First, we queried the RepeatMasker database^36^ and filtered for “Low Complexity”, “Simple Repeat”, and “Satellite” classes. Next, we queried Tandem Repeat Finder (TRF)^37^ to obtain tandem repeats. Finally, we found homopolymers as well as exact repeating sequences with motif size up to 4 bases using a custom script, because the other resources missed some shorter homopolymers and tandem repeats associated with sequencing errors. We identified both perfect repeats (i.e., the same base or bases repeated identically) and imperfect repeats (i.e., repeated sequences with small differences from the repeat motif). To generate the final bed files, we merged these data sources using bedtools (since many regions are expected to overlap) and binned each bed file according to length, since longer repeated regions are expected to be more difficult from a sequencing and variant-calling perspective.

##### Chromosome XY specific regions

Some regions specific to sex chromosomes need to be defined as stratifications due to their importance and distinct evolution of repeats. The human X and Y chromosomes share two pseudoautosomal regions (PARs including PAR1 and PAR2) at the ends of the chromosomes that continue to undergo homologous X-Y recombination^12^. In addition to these, the X-chromosome-transposed region (XTR, sometimes also called PAR3) was duplicated from the X to the Y chromosome in humans after human-chimpanzee divergence and is known to be a recombination hotspot resulting in deletions and inversions^38^. Chromosome Y also includes the ampliconic gene families with highly homologous genes. A typical human Y chromosome harbors 16 single-copy protein-coding X-degenerate genes, with housekeeping functions and homologs on the X chromosome; and 9 protein-coding ampliconic gene families, which have expanded specifically on the Y chromosome^12,38^. Individual stratification BED files were created for PAR, XTR, and ampliconic regions.

##### Patterns of local ancestry in the reference genome

It can be important to know the ancestry of the individuals’ haplotypes that are part of a reference genome because abrupt changes in ancestry of reference regions can cause challenges with linkage disequilibrium (LD) when aligning reads to the reference^39^, and because regions of different ancestry can impact the number of variants called^40^. Local ancestry describes the origin of a chromosomal segment in terms of geographic and regional population. The majority continental super-population affiliation of 1000 Genomes Project samples that most closely match GRCh38 intervals were reported^41^. This was done for African ancestry (AFR), American ancestry (AMR), East-Asian ancestry (EAS), European ancestry (EUR), South-Asian ancestry (SAS). We also report intervals of putative Neanderthal-introgressed origin, based on inferred patterns of identity-by-descent with the Vindija Neanderthal genome ^11,39^. This stratification is currently provided only for GRCh38.

##### Other difficult genomic regions

Some regions of the genome are difficult to analyze due to high degrees of polymorphism or limitations in the reference. The list of “Other difficult genomic regions” includes contigs in the reference assembly that are smaller than 500kb and all gaps in the reference assembly. This category also includes the Major Histocompatibility Complex (MHC) on chromosome 6^42^, the variable/diversity/joining (VDJ) regions on 2, 14 and 22, and the Killer-cell immunoglobulin-like receptor (KIR) region^15^. These three regions are all highly polymorphic and underpin key immunological functions: the MHC region contains the Human Leukocyte Antigen (HLA) genes which determine “donor matches”, the VDJ regions are randomly recombined to produce the T and B cell receptors, and the KIR region codes for one of the key effector receptors on natural killer cells. Stratifications for these difficult regions exist for GRCh37 and GRCh38.

##### Functional, technically difficult to sequence

Several coding regions of the genome present with one of several difficulties. First, some transcription start sites or first exons in the human genome tend to have poor coverage, making accurate sequencing difficult^30^. The stratification defined for these regions include the first 1000 promoters with the lowest coverage by Illumina^30^. Second, some genes are known to be duplicated in most individuals relative to the existing references, which leads to mapping difficulties; this includes KMT2C in both GRCh37 and GRCh38. Third, some genes in the reference are falsely duplicated, which similarly leads to mapping issues. Genes in this category include MRC1 and part of CNR2 (GRCh37) as well as CBS, CRYAA, KCNE1, and H19 (GRCh38).

##### Genome specific

Seven GIAB samples including HG001, HG002, HG003, HG004, HG005, HG006, and HG007 are studied to identify genome-specific difficult regions. These regions cover putative compound heterozygous variants, multiple variants within 50bp of each other, and potential structural variations and copy number variations. These stratifications could be used with benchmarking tools like hap.py to stratify variant calls in terms of true positive, false positive, and false negative.

## Supplementary figures

**Supplementary figure 1.**
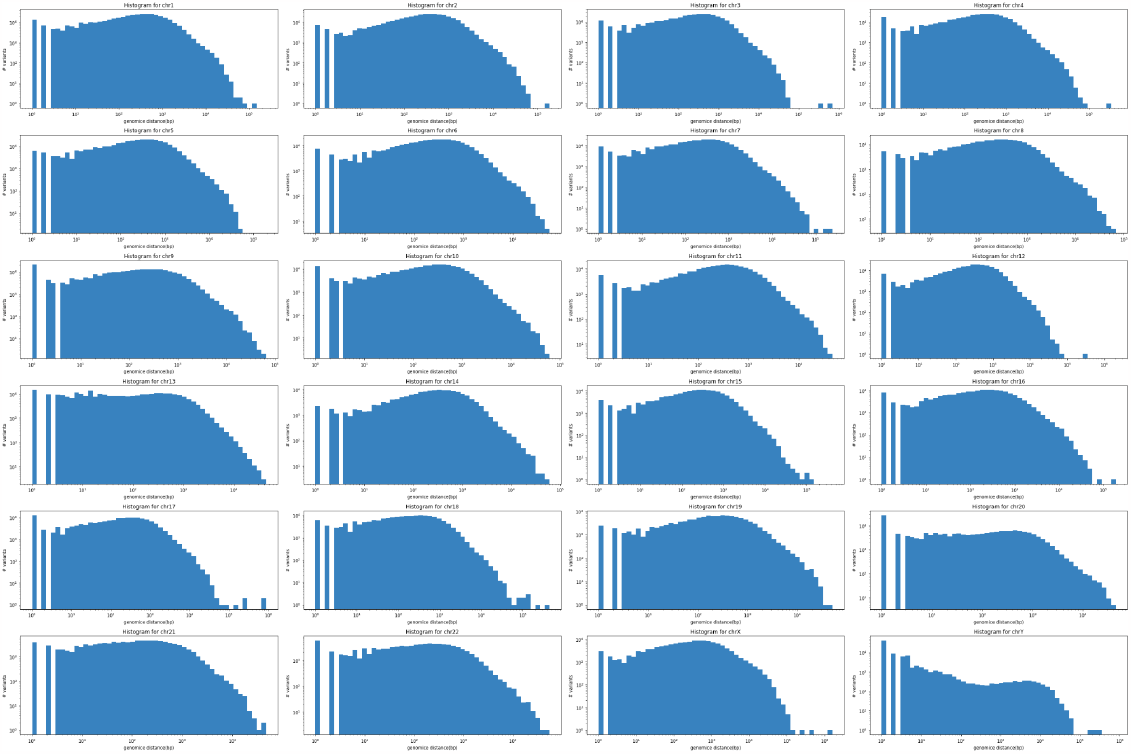
The distribution of genomic distance variants along each chromosome. We should note that a portion of the genome is unknown (existing as Ns in the reference file) and is excluded in this figure. The reason is that no variant can be found in these regions. The distribution of variants along each chromosome is illustrated in Supplementary figure 2. Interestingly, the maximum distance between two consecutive variants (considering N regions) among all chromosomes is 18,014,299 happening between two SNVs at 125,170,338 and 143,184,637 located at 1q12 because this centromeric region is missing from GRCh38.

**Supplementary figure 2.**
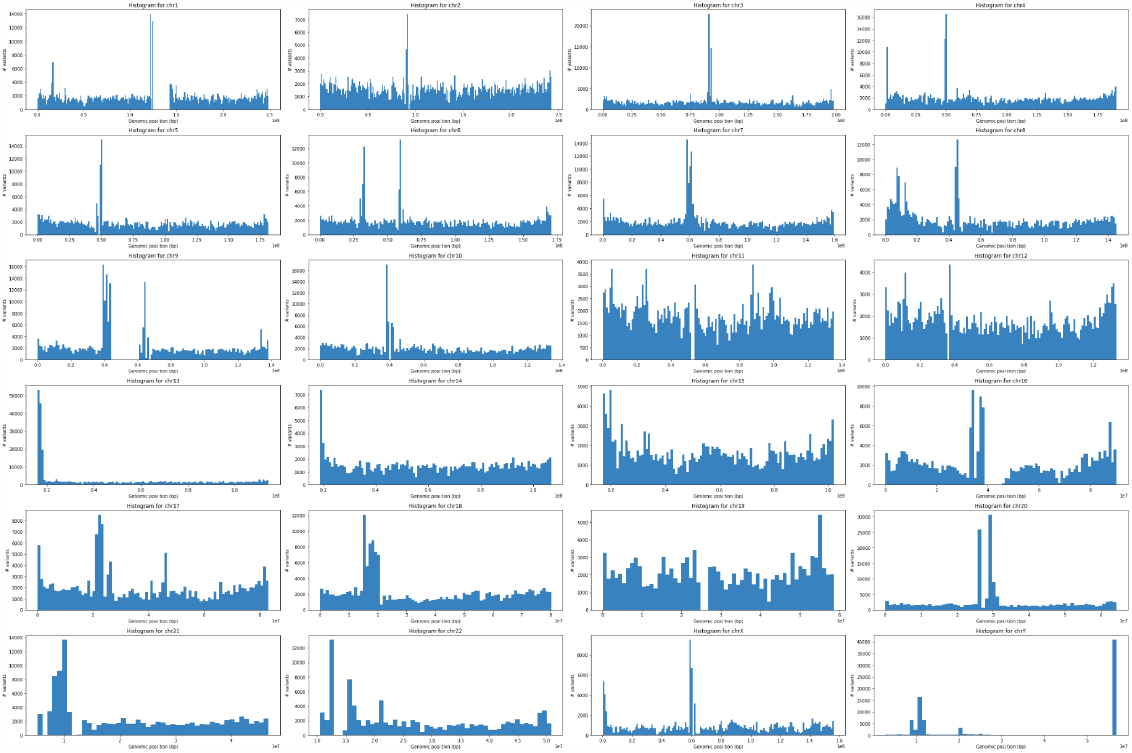
The distribution of variants along each chromosome. This shows how variants are distributed. Y-axis is the number of variants and x-axis is the genomic position of each chromosome.

## References

1. Xiao, C., Zook, J., Trask, S. & Sherry, S. Abstract 5328: GIAB: Genome reference material development resources for clinical sequencing. Cancer Res. 74, 5328–5328 (2014).

2. Wagner, J. et al. Benchmarking challenging small variants with linked and long reads. Cell Genom 2, (2022).

3. Zook, J. M. et al. Integrating human sequence data sets provides a resource of benchmark SNP and indel genotype calls. Nat. Biotechnol. 32, 246–251 (2014).

4. Wagner, J. et al. Curated variation benchmarks for challenging medically relevant autosomal genes. Nat. Biotechnol. 40, 672–680 (2022).

5. Olson, N. D. et al. Variant calling and benchmarking in an era of complete human genome sequences. Nat. Rev. Genet. 24, 464–483 (2023).

6. Sedlazeck, F. J., Lee, H., Darby, C. A. & Schatz, M. C. Piercing the dark matter: bioinformatics of long-range sequencing and mapping. Nat. Rev. Genet. 19, 329–346 (2018).

7. Krusche, P. et al. Best practices for benchmarking germline small-variant calls in human genomes. Nat. Biotechnol. 37, 555–560 (2019).

8. Olson, N. D. PrecisionFDA Truth Challenge V2: Calling variants from short and long reads in difficult-to-map regions. Cell Genomics 2, 100129 (2022).

9. O’Leary, N. A. et al. Reference sequence (RefSeq) database at NCBI: current status, taxonomic expansion, and functional annotation. Nucleic Acids Res. 44, D733–45 (2016).

10. Roy, S. et al. Standards and guidelines for validating next-generation sequencing bioinformatics pipelines: A joint recommendation of the Association for Molecular Pathology and the College of American Pathologists. J. Mol. Diagn. 20, 4–27 (2018).

11. Nurk, S. et al. The complete sequence of a human genome. Science 376, 44–53 (2022).

12. Rhie, A. et al. The complete sequence of a human Y chromosome. bioRxiv 2022.12.01.518724 (2022) doi:10.1101/2022.12.01.518724.

13. Antonarakis, S. E. Short arms of human acrocentric chromosomes and the completion of the human genome sequence. Genome Res. 32, 599–607 (2022).

14. Foox, J. et al. Performance assessment of DNA sequencing platforms in the ABRF Next-Generation Sequencing Study. Nat. Biotechnol. 39, 1129–1140 (2021).

15. Pyke, R. M. et al. Computational KIR copy number discovery reveals interaction between inhibitory receptor burden and survival. Pac. Symp. Biocomput. 24, 148–159 (2019).

16. Jarvis, E. D. et al. Semi-automated assembly of high-quality diploid human reference genomes. Nature 611, 519–531 (2022).

17. Smolka, M., Rescheneder, P., Schatz, M. C., von Haeseler, A. & Sedlazeck, F. J. Teaser: Individualized benchmarking and optimization of read mapping results for NGS data. Genome Biol. 16, 235 (2015).

18. Chen, N.-C., Solomon, B., Mun, T., Iyer, S. & Langmead, B. Reference flow: reducing reference bias using multiple population genomes. Genome Biol. 22, 8 (2021).

19. Majidian, S., Kahaei, M. H. & de Ridder, D. Hap10: reconstructing accurate and long polyploid haplotypes using linked reads. BMC Bioinformatics 21, 253 (2020).

20. Chin, C.-S. et al. Phased diploid genome assembly with single-molecule real-time sequencing. Nat. Methods 13, 1050–1054 (2016).

21. Dwarshuis, N. et al. StratoMod: Predicting sequencing and variant calling errors with interpretable machine learning. bioRxiv 2023.01.20.524401 (2023) doi:10.1101/2023.01.20.524401.

22. Pedersen, B. S. et al. Effective variant filtering and expected candidate variant yield in studies of rare human disease. NPJ Genom Med 6, 60 (2021).

23. Majidian, S. & Sedlazeck, F. J. PhaseME: Automatic rapid assessment of phasing quality and phasing improvement. Gigascience 9, (2020).

24. Yip, K. Y., Cheng, C. & Gerstein, M. Machine learning and genome annotation: a match meant to be? Genome Biol. 14, 205 (2013).

25. Fotsing, S. F. et al. The impact of short tandem repeat variation on gene expression. Nat. Genet. 51, 1652–1659 (2019).

26. Turner, S. et al. Quality control procedures for genome-wide association studies. Curr. Protoc. Hum. Genet. Chapter 1, Unit1.19 (2011).

27. Rautiainen, M. et al. Telomere-to-telomere assembly of diploid chromosomes with Verkko. Nat. Biotechnol. 41, 1474–1482 (2023).

28. Derrien, T. et al. Fast computation and applications of genome mappability. PLoS One 7, e30377 (2012).

29. Baid, G. et al. An Extensive Sequence Dataset of Gold-Standard Samples for Benchmarking and Development. bioRxiv 2020.12.11.422022 (2020) doi:10.1101/2020.12.11.422022.

30. Ross, M. G. et al. Characterizing and measuring bias in sequence data. Genome Biol. 14, R51 (2013).

31. Zhao, S., Agafonov, O., Azab, A., Stokowy, T. & Hovig, E. Accuracy and efficiency of germline variant calling pipelines for human genome data. Sci. Rep. 10, 20222 (2020).

32. Li, H. seqtk: Toolkit for processing sequences in FASTA/Q formats. (Github, 2023).

33. Vollger, M. R. et al. Segmental duplications and their variation in a complete human genome. Science 376, eabj6965 (2022).

34. Vollger, M. R. et al. Long-read sequence and assembly of segmental duplications. Nat. Methods 16, 88–94 (2019).

35. Bailey, J. A., Yavor, A. M., Massa, H. F., Trask, B. J. & Eichler, E. E. Segmental duplications: organization and impact within the current human genome project assembly. Genome Res. 11, 1005–1017 (2001).

36. RepeatMasker website. http://www.repeatmasker.org (2023).

37. Benson, G. Tandem repeats finder: a program to analyze DNA sequences. Nucleic Acids Res. 27, 573–580 (1999).

38. Cotter, D. J., Brotman, S. M. & Wilson Sayres, M. A. Genetic Diversity on the Human X Chromosome Does Not Support a Strict Pseudoautosomal Boundary. Genetics 203, 485–492 (2016).

39. Aganezov, S. et al. A complete reference genome improves analysis of human genetic variation. Science 376, eabl3533 (2022).

40. Goetz, L. H., Uribe-Bruce, L., Quarless, D., Libiger, O. & Schork, N. J. Admixture and clinical phenotypic variation. Hum. Hered. 77, 73–86 (2014).

41. Lowy, E., Fairley, S. & Flicek, P. Variant calling across 505 openly consented samples from four Gambian populations on GRCh38. Wellcome Open Res. 6, 239 (2021).

42. Chin, C.-S. et al. A diploid assembly-based benchmark for variants in the major histocompatibility complex. Nat. Commun. 11, 4794 (2020).

